# Drought reduces formation, but enhances persistence, of mineral-associated organic matter in a grassland soil

**DOI:** 10.1101/2025.05.23.655838

**Authors:** Noah W. Sokol, Megan M. Foley, Steven J. Blazewicz, Nicole DiDonato, Katerina Estera-Molina, Mary Firestone, Alex Greenlon, Bruce A. Hungate, William Kew, Ljiljana Paša-Tolić, Eric Slessarev, Jennifer Pett-Ridge

## Abstract

Drought effects are pervasive in many terrestrial ecosystems, yet little is known about the impact of drought on the transformation of plant C inputs to mineral-associated organic matter (MAOM) – the largest and slowest-cycling pool of organic carbon (C) on land. Using ^13^C-CO_2_ greenhouse labeling chambers, we tracked the formation of ^13^C-MAOM derived from *Avena barbata* living root inputs (^13^C-rhizodeposits) versus *A. barbata* decaying root inputs (^13^C-root detritus) under normal moisture and spring drought conditions in a semi-arid grassland soil, and then tested the durability of this ^13^C-MAOM in a subsequent persistence assay. Overall, drought reduced formation of MAOM – both per gram of soil and across the entire soil profile. Notably, drought conditions enhanced the persistence of MAOM derived from root detritus, though not of MAOM derived from rhizodeposition. Drought had the most pronounced effect on MAOM accrual from rhizodeposition late in plant development (week 12) whereas it had the most pronounced effect on MAOM accrual from root detritus early in root decomposition (week 4). These temporal responses were associated with distinct trajectories in microbial community-level growth rates, the average mass of OM compounds, and the number of unique metabolites within each habitat. Our results provide mechanistic evidence that drought reduces overall formation of MAOM but can enhance its persistence in a grassland soil.

## INTRODUCTION

Mineral-associated organic matter (MAOM) is Earth’s largest and slowest-cycling pool of terrestrial organic carbon (C) (Heckman et al. 2022). MAOM-C is primarily derived from plant C inputs entering the mineral soil, which are subsequently transformed to MAOM via the activities of soil microbes (Sokol et al. 2019, 2022). The transformation of these plant C inputs into MAOM draws down atmospheric CO_2_ into soil, which can persist over long time scales, though a portion of MAOM can cycle rapidly (Lavallee et al. 2020; Jilling et al. 2025). Yet, little is known about how MAOM formation and persistence will respond to changes in precipitation – especially in grasslands (Schimel 2018; Jilling et al. 2024). Grasslands are particularly susceptible to increasing drought events (Wu et al. 2022), and more than 70% of soil organic C in grasslands exists as MAOM-C (Sokol et al. 2022). Soil moisture influences MAOM cycling through its effects on net primary productivity (NPP), soil microbial growth and activity, and the movement of solutes through the soil matrix (Churkina et al. 1999; Schimel 2018). The abundance and persistence of MAOM is strongly linked to moisture availability (Heckman et al. 2023), yet few manipulative studies have directly tested how drought affects the transformation of plant C inputs into MAOM and its subsequent persistence (Canarini and Dijkstra 2015; Canarini et al. 2017; Jilling et al. 2024).

There are contrasting hypotheses for how drought may affect MAOM (Schimel 2018). Drought may decrease MAOM formation by decreasing net primary productivity (NPP) and plant C inputs to the soil, especially belowground root inputs, which are the dominant source of plant C to the MAOM pool (Rasse et al. 2005; Sokol et al. 2019, 2022; Zhang and Xi 2021). Drought may reduce microbial activity and growth, thus decreasing the microbial transformation of plant C compounds to MAOM (Hueso et al. 2012; Fuchslueger et al. 2019; Domeignoz-Horta et al. 2020). Moisture limitation can also decrease diffusion of compounds through the mineral matrix, reducing interactions between OM and minerals. Alternatively, drought may increase MAOM by enhancing interactions between soluble organic compounds and mineral surfaces through increased water tension during dry down events (Kaiser et al. 2015), or by stimulating the synthesis of specific root and microbial products, such as extracellular polymeric substances – which are posited to be dominant precursors of MAOM (Sher et al. 2020; Benard et al. 2023; Sokol et al. 2024).

Drought may have similar or different effects on MAOM formation versus MAOM persistence – i.e., how much new MAOM is formed versus the proportion of that MAOM that persists over a defined timeframe. For instance, a recent observational field study of 34 field sites in the conterminous United States found that MAOM abundance (MAOM-C %) and persistence (measured via the radiocarbon age of MAOM-C, Δ^14^C (‰)) were negatively related in humid soils, though there was no relationship between MAOM abundance and persistence in arid soils (Heckman et al. 2023). This observation was posited to be the result of higher NPP in humid environments, and greater abundance and biochemical quality of root inputs into the soil (e.g., low molecular weight root exudates). These factors contributed to a large standing stock of MAOM in humid environments, but also to one that was more susceptible to priming and decomposition, and thus cycled more quickly than MAOM in arid environments (Heckman et al. 2023). While compelling, it is unclear how these hypotheses manifest within an individual ecosystem experiencing drought. There is a clear need for mechanistic studies that directly test how drought affects the formation and persistence of MAOM – especially in ecosystems affected by altered precipitation regimes (Sokol et al. 2022; Heckman et al. 2023).

In grasslands, MAOM is primarily derived from two sources of root input. Living root inputs (rhizodeposits) generate a microbial habitat around the living root (the rhizosphere), whereas decaying root inputs form a microbial habitat around root detritus (the detritusphere) (Pett-Ridge et al. 2021; Sokol et al. 2024; Witzgall et al. 2024). Key differences between these two sources of plant C may lead to distinctive impacts of drought on their transformation to MAOM (Sokol et al. 2024). Rhizodeposition – a mix of lower molecular weight exudates (sugars, proteins, organic acids), as well as higher molecular weight compounds, such as mucilage and border cells – can change in amount and composition over the course of a growing season and in response to environmental conditions like drought (Naylor and Coleman-Derr 2018; Williams and de Vries 2020). While drought can impact the total amount of root biomass produced (Guasconi et al. 2023), decaying root detritus is not as dynamically responsive as rhizodeposition to environmental conditions. Rather, decomposition of root litter should follow a more predictable trajectory (Poll et al. 2008), which may be delayed under drought conditions.

Here, in a manipulative ^13^C-tracer greenhouse experiment, we directly tested how drought affected the formation and short-term persistence of MAOM derived from belowground plant inputs in a semi-arid grassland soil. Over a 12-week growing period, we tracked ^13^C- rhizodeposition from living *A. barbata* plants versus decaying *A. barbata* root litter into soil organic matter (SOM) pools, under both normal moisture versus drought conditions. We also assessed the relative persistence of ^13^C-MAOM that was formed at the end of 12-week period under normal moisture versus drought conditions, via a subsequent 90-day persistence assay (Kallenbach et al. 2016; Oldfield et al. 2018; Whalen et al. 2024). By pairing data on ^13^C- MAOM accrual with microbial community-level growth rate, as well as the chemical composition of SOM (measured via FTICR-MS) and lipid and metabolite composition in the rhizosphere and detritusphere, we asked: (1) How does drought affect the accrual and chemical composition of MAOM derived from living versus dead root inputs over the course of a growing season? (2) Do MAOM formation and MAOM persistence respond the same way to drought in different microbial habitats? (3) What plant and microbial dynamics help explain these patterns?

## METHODS

### Experimental Design

To test how a spring drought affects the formation and persistence of MAOM in a grassland soil, we conducted a ^13^C-labeling greenhouse tracer and soil moisture manipulation study for a 12- week period. The experimental design is described in detail in Sokol et al. (2024). Briefly, soil was collected at 0-10 cm depth (‘A’ mineral horizon) below a stand of *A. barbata* at the University of California Hopland Research and Extension Center (39°00.106‵N, 123°04.184‵W) – a Mediterranean annual grassland ecosystem (MAT max/min = 23/7 C; MAP = 956 mm yr^-1^), where *Avena* spp. is the dominant vegetation. The soil is classified as a Typic Haploxeralf, has a pH of ∼5.6, and contains 45% sand, 36% silt, and 19% clay, and initial C and N content of 2.2% and 0.24%, respectively.

After soil collection, roots were removed, and soil was passed through a 2-mm sieve and packed to field bulk density (1.21 g cm^-3^) in rectangular acrylic microcosms (11.5 × 2.9 × 25.5 linear cm). Two types of microcosms were established: planted microcosms (‘rhizodeposit’ treatment), where ^13^C-CO_2_ was tracked through living *A. barbata* plants into the rhizosphere, and unplanted microcosms (the ‘root detritus’ treatment), where ^13^C-labeled *A. barbata* root detritus was tracked into the detritusphere (Fig. S1). In unplanted microcosms, a 28-μm mesh bag was buried in the microcosm center, which contained ∼65 g of Hopland soil mixed with 1-5 mm fragments of either ^13^C-labeled *A. barbata* root detritus (77 ± 1.7 atom% ^13^C-labeled), or natural abundance *A. barbata* root detritus, to a concentration of 0.013 g root detritus dry g soil^-1^. *Avena barbata* root detritus was generated using the same source of seeds used in the main experiment, and plants were grown for 7 weeks in a mix of Hopland soil and sand immediately prior to the experimental period. After 2 weeks of plant establishment, half the plants were exposed for 5 weeks of continuous ^13^CO_2_ labeling (same protocol as described below), and the other half were grown in natural abundance environment. After 7 weeks of growth, roots were harvested, washed, dried to constant temperature, and cut into 1-5 mm fragments, before being packed into detritusphere mesh bags.

All unplanted microcosms were incubated in growth chambers with a natural abundance CO_2_ headspace throughout the experimental period. In planted microcosms, three germinated *A. barbata* seedlings were planted evenly apart in the soil, and microcosms were then inside growth chambers (56 × 56 × 76 linear cm) at the Environmental Plant Isotope Chamber (EPIC) facility at the Oxford Tract Greenhouse at University of California Berkeley (Pett-Ridge and Firestone 2017; Pett-Ridge et al. 2021). After a two-week seeding establishment phase, a subset of the planted microcosms (*n* = 6 per timepoint) were grown inside ^13^CO_2_ growth chambers for 10 continuous weeks, which were filled with ∼99 atom% ^13^CO_2_ (Sigma Aldrich) and a CO_2_ setpoint of 400 ppm. A control set of planted microcosms (*n* = 4) were grown in natural abundance (^12^CO_2_) growth chambers, and used for measurements that did not require a ^13^C-label – including FTICR-MS, lipidomics, and metabolomics (discussed further below). Full details on chamber maintenance and monitoring (including temperature, moisture, light, CO_2_ concentration ^13^C enrichment, and other parameters) are fully described in Sokol et al. (2024).

Planted and unplanted microcosms were maintained at one of two moisture treatments during the 12-week experimental period: ‘normal moisture’ (∼16% ± 0.3 gravimetric soil moisture; mean ± standard error) or ‘spring drought’ conditions (∼8% ± 0.5 gravimetric soil moisture). These treatments simulated differences in soil moisture during the spring growing season in California semiarid grasslands. To monitor soil moisture, all microcosms were weighed twice weekly, and adjusted to the correct soil moisture by mass (Shi et al. 2015). At each harvest (4, 8, and 12 weeks), gravimetric soil moisture was measured for all microcosms. As described in Sokol et al. (2024), it was confirmed that the hydrological conditions of the detritusphere mesh bags were not significantly different from the rhizosphere.

### Sample collection

Aboveground and belowground plant biomass in planted microcosms was collected at each harvest. Aboveground *A. barbata* biomass was clipped at the base of the stems, dried at 65°C and weighed. Rhizosphere soil was collected by gently shaking the root systems to remove loosely attached soil; rhizosphere soil was defined as soil still clinging to the roots (Nuccio et al. 2020). The roots were washed, dried at 65°C and weighed. A subset of rhizosphere soil was immediately placed within a 15-mL falcon tube on dry ice and stored at -80°C. The remaining rhizosphere soil was carefully separated from the roots by hand; a subset was kept fresh at room temperature for measuring microbial community-level growth rate and other assays (described below), and a subset was air-dried for SOC analysis. In unplanted microcosms detritusphere soil was collected from inside the 28-μm detritusphere mesh bag, avoiding any fragments of decaying root material. Subsets of detritusphere soil were stored at -80°C, fresh, and air-dried.

### SOC pools

We used a combined density and physical fractionation protocol to measure three distinct SOC fractions – mineral-associated organic matter (MAOM), particulate organic matter (POM), and coarse heavy associated organic matter (CHAOM) (Leuthold et al. 2024; Sokol et al. 2024). First, 25 mL of 1.85 g cm^3^ sodium polytungstate (SPT) was added to 5 g of air-dried soil in a 50-mL falcon tube and shaken on a reciprocal shaker (∼200 oscillations/minute) for 18 hours with glass beads. The supernatant was filtered with a glass fiber filter to separate the light particulate fraction (<1.85 g cm^-3^) – defined as POM. The pelletized heavy fraction at the base of the tube (<1.85 g cm^-3^) was rinsed repeatedly to remove excess SPT, vortexed with 25-mL deionized water and passed through a 53-µm sieve. The < 53-µm fraction (clay + fine silt) is defined as MAOM; the > 53-µm fraction is defined as CHAOM (Soong and Cotrufo 2015; Leuthold et al. 2024). All samples were dried, ground, weighed, and analyzed for % total C and δ^13^C on an elemental analyzer coupled to an isotope ratio mass spectrometer (EA-IRMS; Costech ECS 4010, Costech Analytical Technologies, Valencia, CA, USA).

Atom% ^13^C enrichment of each SOM pool was calculated by subtracting atom% ^13^C of mineral-associated SOC at *t* = 0 from the atom% ^13^C in the enriched sample. The total µg of ^13^C that accumulated in each SOM pool (i.e., MAOM, POM, CHAOM) were then calculated using a standard mixing model (Kallenbach et al. 2016; Sokol et al. 2024).

### SOC persistence

To measure the short-term persistence of the ^13^C-MAOM fraction, we exposed a subsample of each ^13^C-labeled rhizosphere or detritusphere from week 12 to a subsequent 90- day persistence assay (Bradford et al. 2013; Kallenbach et al. 2016; Oldfield et al. 2018; Whalen et al. 2024). During this 90-day period, soils were incubated at 30°C and 65% water holding capacity, and frequently hand-mixed, to provide ideal conditions for microbial activity such that all easily available C is mineralized (Bradford et al. 2013). The concentration of ^13^C-MAOM g soil^-1^ remaining at the end of the persistence assay relative to the concentration of ^13^C-MAOM at the beginning of the persistence assay is considered a proxy of more persistent ^13^C-MAOM (Kallenbach et al. 2016; Oldfield et al. 2018; Whalen et al. 2024). The concentration of ^13^C- MAOM (µg ^13^C g soil^-1^) at the end of the persistence assay was calculated using a standard isotope mixing model. Then, the concentration of ^13^C-MAOM at the end of the assay was divided by the concentration of ^13^C-MAOM in that sample at the beginning of the assay and multiplied by 100%, to give a % of ^13^C-MAOM that remained and was considered more persistent.

### Microbial community-level growth rate

To measure microbial community-level growth rate, we used the ^18^O-H_2_O method, as described by Spohn et al (2016). Applying this method to capture microbial growth rates in normal moisture and drought treatments in this particular soil is described in detail in Sokol et al. (2024). Briefly, a 1-g sample of fresh soil was weighed into a 20-mL Wheaton glass serum vial, and a combination of ^18^O-H_2_O (98 atom % H_2_^18^O, Isoflex, San Francisco, CA, USA) and natural abundance H_2_O was added to the sample so the resulting soil water solution was ∼20 atom% ^18^O. Vials were capped, incubated for 72-hours at room temperature, then 5-mL of gas from the vial headspace was collected for CO_2_ analysis (and respiration rate) on a gas chromatograph (Agilent 7890B GC System). After sampling the headspace, the soil was removed and immediately flash frozen in liquid nitrogen and stored at - 80°C. DNA was extracted from frozen soil using a Qiagen DNEasy PowerSoil Pro kit following the manufacturer’s instructions; the bead beating step was performed for 1 minute at 5.5 m^2^/s on a FastPrep. DNA was quantified using the Qubit DNA BR Assay Kit (ThermoFischer Scientific). A 50 μL aliquot of DNA was dried at 60°C in a pre-weighed silver capsule spiked with 100 μL salmon sperm DNA (to achieve the oxygen detection limit) and analyzed for δ^18^O and total O content (μg) on a Thermochemical Element Analyzer (TC/EA) coupled to an IRMS.

Microbial community-level growth rate was calculated from the amount of new DNA produced during the incubation period (tracked by ^18^O incorporation into microbial DNA during growth). The amount of new DNA produced during the incubation period (DNA_p_) is the difference in ^18^O abundance between DNA from labeled and control incubations times the proportion (by mass) of DNA that is oxygen (0.3121) divided by the length of the incubation and mass of soil incubated. The conversion mass ratios of MBC:DNA for each sample was applied to calculate growth rate (Gr) in mg C day^-1^ g^-1^ soil:

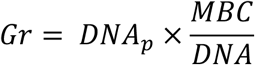

Where DNA_p_ (µg DNA day^-1^ g^-1^ dry soil) is the DNA produced during the incubation, MBC (mg C g^-1^ dry soil) is microbial biomass C measured via chloroform fumigation extraction (described in Sokol et al., 2024), and DNA is the soil DNA concentration (µg DNA g^-1^ dry soil) determined via Qubit.

### Fourier transform ion cyclotron resonance mass spectrometry (FTICR-MS), Metabolomics, Lipidomics

Natural abundance samples were used for FTICR-MS, metabolite and lipid analyses. On each sample, a water extraction was first performed for FTICR-MS analysis, followed by a modified Folch type extraction which yielded polar and non-polar liquid fractions containing metabolite and lipid components. Full details of the extractions, analysis, and data processing are included in Supporting Information (Supp. Text 1).

### Statistical analysis

All analyses were conducted in R version 4.2.1 (R Core Team, 2022). We used multiple regression to analyze the effect of soil moisture and time on ^13^C-SOC pools (MAOM, POM, CHAOM), aboveground and belowground plant biomass, microbial community-level growth rate, amino sugars, proteins, and carbohydrates, the total mass of OM compounds, and the total number of metabolites. We used simple regression to test for the effect of moisture on persistence of MAOM. Multiple and simple regression models were run separately for the rhizosphere and the detritusphere. All models were screened for normality of residuals (Shapiro-Wilk test) and heteroscedasticity of residuals (visual assessment of residual plots).

For metabolites, as most of the detected features were not annotated identities, an analysis of all the detected features was performed. Briefly, data across polarities and modalities were collated, blank subtracted (blank sample median signal + 3 standard deviations threshold), and retaining only features detected in 3 out of 4 replicate measurements. Volcano plots were produced to highlight the metabolite features detected to have significant (ANOVA, p<0.05) and log2 fold differences (log2FC > 1 or <-1) in abundances between conditions for each timepoint and habitat. For lipids, at timepoints 1 and 3, the positive and negative lipid data were combined and statistically significant lipid abundances identified through ANOVA analysis. Tukey’s HSD was performed on significant (p<0.05) lipids to validate significance. The 25 most significantly different lipids were visualized as a heatmap of their log-normalized abundance for each sample.

## RESULTS

### Soil organic matter and plant biomass

During the 12-week spring growing season, drought reduced formation of ^13^C-MAOM in the rhizosphere and detritusphere – both the concentration of ^13^C-MAOM per g of soil (µg ^13^C g soil^-1^) and the formation of ^13^C-MAOM at the whole microcosm scale (µg ^13^C microcosm^-1^) (Fig. 1). Drought effects on MAOM showed contrasting temporal patterns in the rhizosphere versus the detritusphere. The most pronounced effect of drought on ^13^C-MAOM formation in the rhizosphere occurred at week 12 – the end of the growing season. This effect at week 12 was observed for both ^13^C-MAOM per g of rhizosphere soil (Fig. 1a; time*moisture interaction, p=0.0003) and across the entire rhizosphere at the whole microcosm scale (Fig. 1b; time*moisture interaction, p<0.0001). In these same microcosms with living plants (i.e., the ‘rhizodeposit-only’ microcosm), drought also had the most pronounced at week 12 on aboveground biomass (moisture*time interaction; p<0.001; Fig. S2a) and total belowground root biomass (moisture*time interaction p = 0.04; Fig. S2b). In contrast with the rhizosphere, drought had the most pronounced effect on detritusphere ^13^C-MAOM at week 4, during the first stages of root decomposition (Fig. 1 c,d). At week 4, there was lower ^13^C-MAOM per g of detritusphere soil (p = 0.02; Fig 1c) and across the entire detritusphere in the whole microcosm (p= 0.01 Fig. 1d). By week 12, there was no difference in detritusphere ^13^C-MAOM per g of soil (p=0.9; Fig. 1c) or within the whole microcosm (p=0.6; Fig. 1d) in normal moisture versus drought conditions.

**Figure 1.**
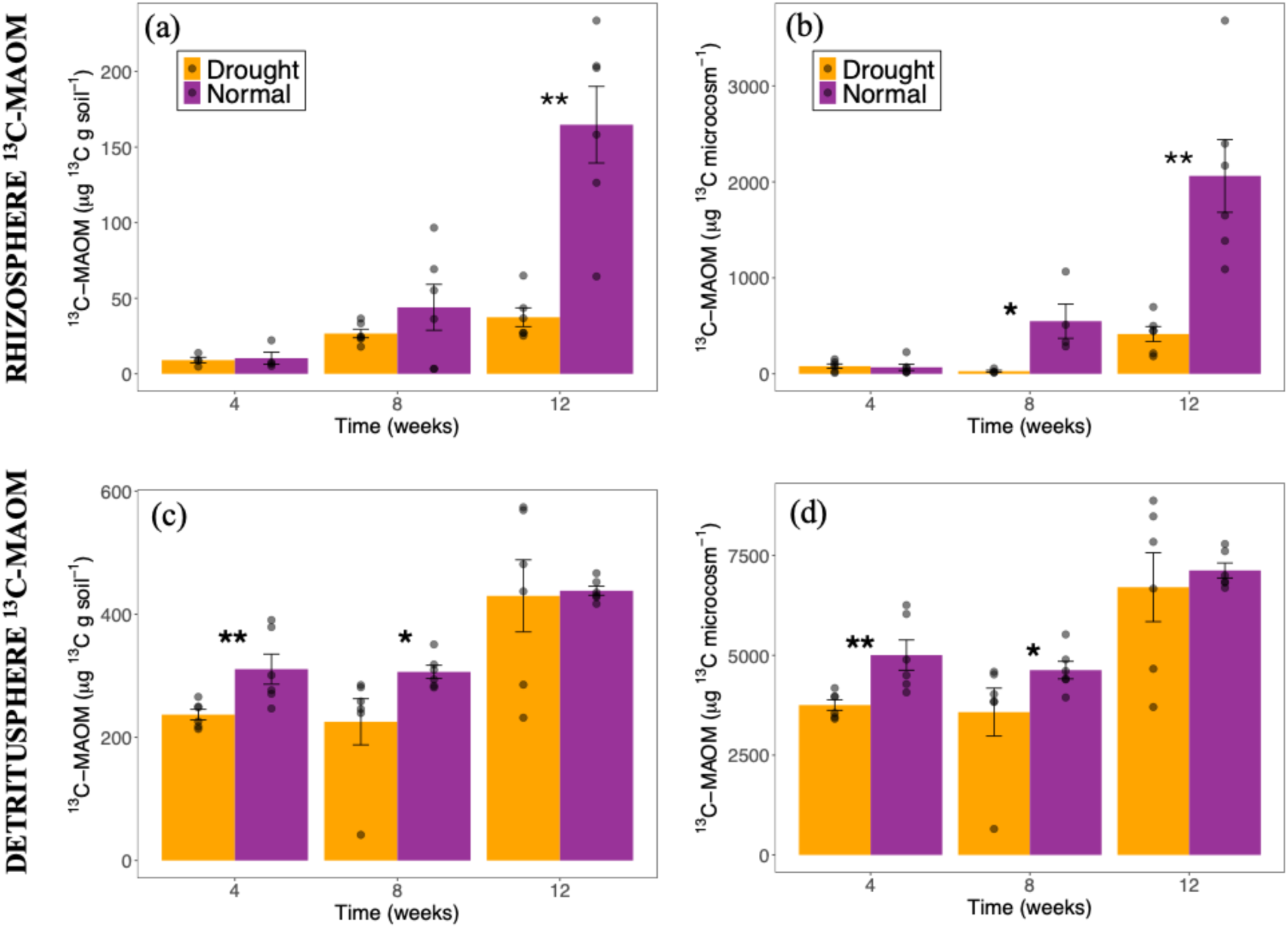
^13^C-mineral associated organic matter (^13^C-MAOM) formed over a 12-week period in a ^13^CO_2_ greenhouse labeling and moisture manipulation experiment in a semi-arid California annual grassland soil. Shown below is ^13^C-MAOM formed in the rhizosphere (a,b) and detritusphere (c,d) of under normal moisture (purple) or droughted conditions (orange). Asterisks indicate significant differences between moisture treatments at a given time point (* indicates p<0.05; ** indicates p<0.005). *N* = 6. Left panels (a,c) show ^13^C-MAOM formation per g soil (ug ^13^C g soil^-1^); right panels show ^13^C-MAOM formation at the whole microcosm scale (ug ^13^C microcosm^-1^).

As with the ^13^C-MAOM pool in the rhizosphere, the ^13^C-POM pool in the rhizosphere increased over the 12-week growing season (Fig. S3a). Drought had the most significant effect on the rhizosphere ^13^C-POM pool at week 12 (moisture*time interaction, p = 0.008; Fig. S3a). Drought also trended towards having the strongest effect on the rhizosphere ^13^C-CHAOM pool at week 12 (moisture*time interaction, p = 0.1; Fig. S3b). In contrast with the rhizosphere, the detritusphere ^13^C-POM pool decreased over the 12-week period (p = 0.03; Fig. S3c). While drought decreased ^13^C-POM formation in the rhizosphere, drought was associated with a larger ^13^C-POM pool in the detritusphere (p = 0.09; Fig. S3d), due to slower decomposition under drought. There was no significant effect of time (p = 0.3) or soil moisture (p = 0.97) on the size of the ^13^C-CHAOM pool in the detritusphere (Fig. S3d).

### *13C-MAOM* persistence assay

In the rhizosphere, there was no difference in the persistence of rhizosphere ^13^C-MAOM that formed under normal moisture versus drought conditions (i.e., the % of ^13^C-MAOM remaining after a 90-day persistence assay) (p= 0.02; Fig. 2a). In the detritusphere, ^13^C-MAOM that formed under drought conditions was more persistent that ^13^C-MAOM that formed under normal moisture (p=0.0006; Fig. 2b).

**Figure 2.**
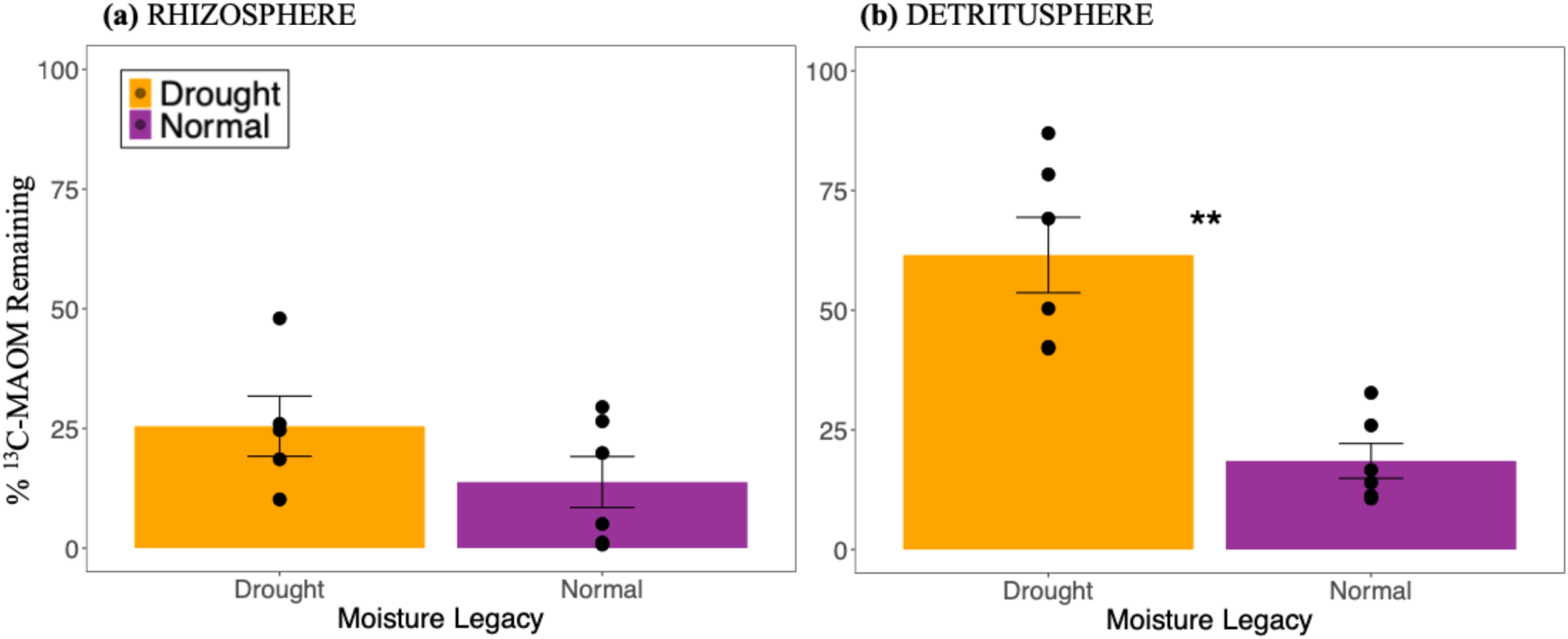
Persistence of ^13^C-mineral associated organic matter (MAOM), measured as the % of ^13^C-MAOM remaining after a 90-day persistence assay after a 12-week ^13^CO_2_ labeling experiment. Purple bars indicate ^13^C-MAOM formed under normal moisture conditions, orange bars indicate ^13^C-MAOM formed under drought conditions. Asterisks indicate significant differences between moisture treatments at a given timepoint (** indicates p<0.005).

### Microbial community-level growth rate

Drought reduced microbial community-level growth rate in the rhizosphere and detritusphere (Fig. 3). As with MAOM, microbial community-level growth rate followed opposing patterns through time in the rhizosphere versus detritusphere. In the rhizosphere, drought had the strongest negative effect at week 12, when microbial community-level growth was at its highest (moisture*time interaction, p = 0.02; Fig. 3a). In the detritusphere, microbial community level growth rate decreased through time (p<0.001) and was lower under drought conditions (p=0.01; Fig. 3b).

**Figure 3.**
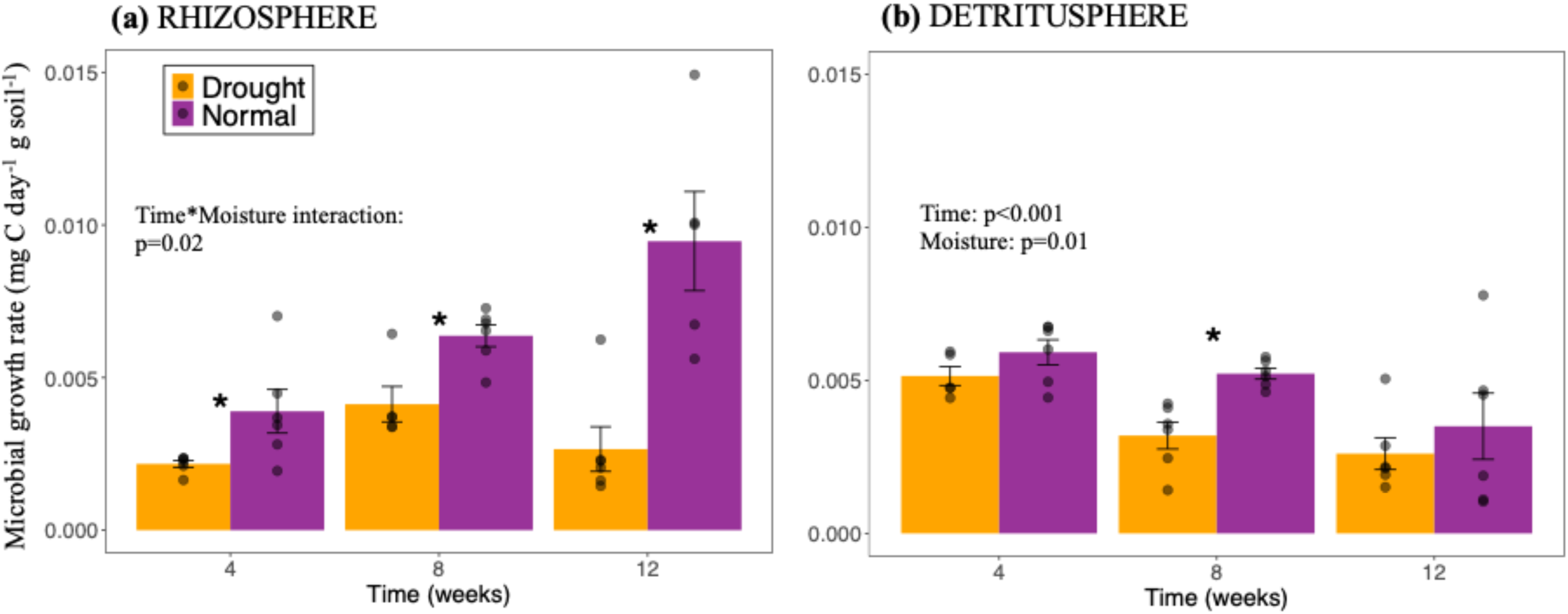
Microbial community-level growth rate in the rhizosphere and detritusphere of a semi-arid grassland soil over a 12-week ^13^CO_2_ labeling experiment. Microbial community-level growth rate (‘microbial growth rate’; mg C day^-1^ soil^-1^) was measured via the ^18^O-H_2_O method (see Methods). Asterisks indicate significant differences between moisture treatments at a given timepoint (* indicates p<0.005). P values indicate if time or moisture treatment (or their interaction) were significant in linear models. *N*=6.

### Chemical composition of organic matter (FTICR-MS)

The chemical composition of soil organic matter changed through time in contrasting ways in the rhizosphere and detritusphere, as measured through FTICR-MS (Fig. 4). In the rhizosphere, amino-sugar like compounds (p < 0.001), protein-like compounds (p = 0.02), and carbohydrate-like compounds (p = 0.001) increased through time over the 12-week growing season (Fig. 4a-c). Carbohydrate-like compounds were lower under drought in the rhizosphere (p = 0.06). In contrast with the rhizosphere, protein-like compounds (p = 0.05) and carbohydrate-like compounds (p = 0.006) decreased through time in the detritusphere. All three compound classes were more abundant under drought relative to normal moisture conditions in the detritusphere (Fig. 4e-g). In contrast, the average mass of OM compounds decreased through time in the rhizosphere (p = 0.0007), and was lower under drought conditions (p = 0.01; Fig. 4d). The average mass of OM compounds increased over time in the detritusphere (p = 0.004) and was higher under normal moisture conditions (p = 0.04; Fig. 4h).

**Figure 4.**
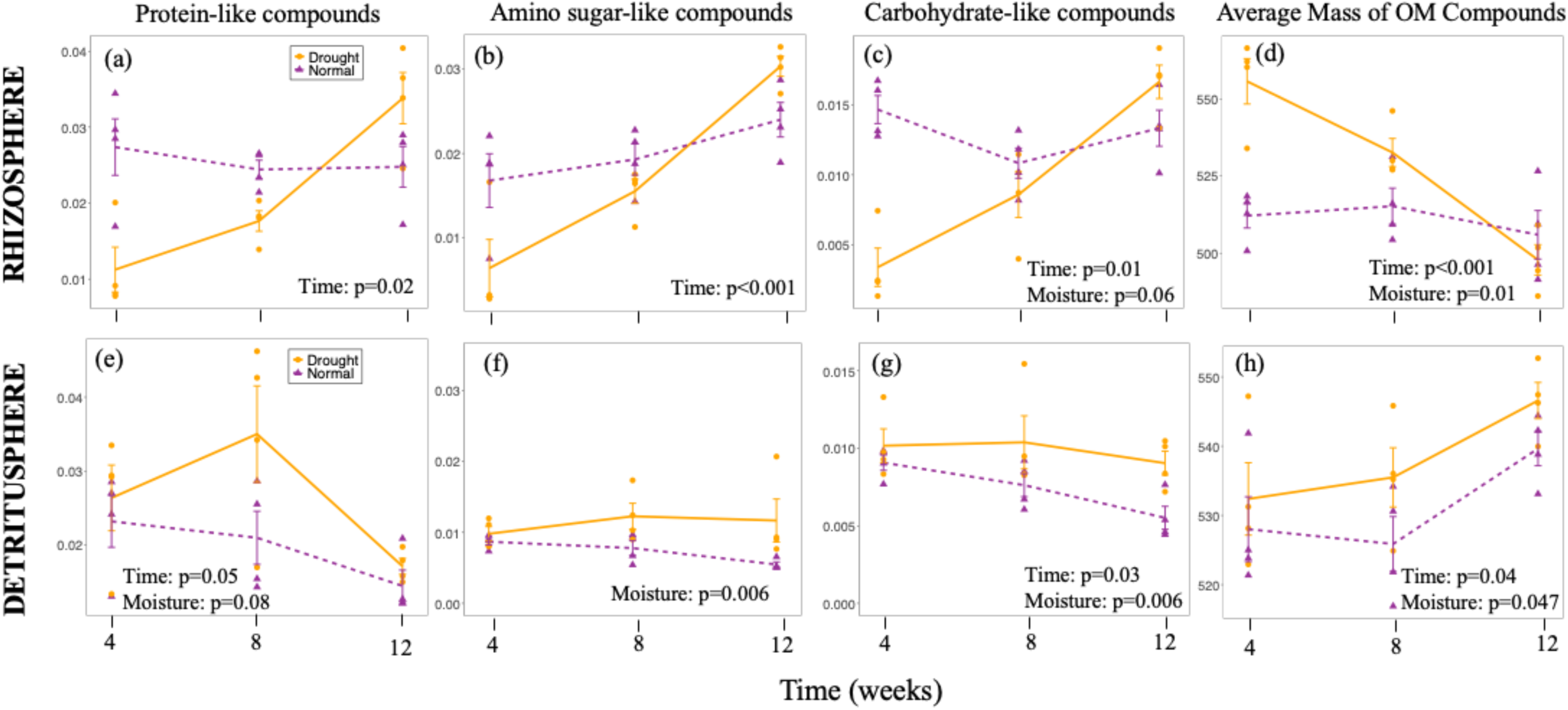
Chemical composition of soil organic matter in the rhizosphere and detritusphere of a semi-arid grassland soil over a 12- week ^13^CO_2_ labeling experiment, measured through FTICR-MS. Normal moisture shown in purple; droughted conditions shown in orange. Upper panels show the rhizosphere (a-d); lower panels show the detritusphere (e-h). The *y*-axis units for protein-like compounds, amino sugar-like compounds, and carbohydrate-like compounds is ‘average % of formulas.’ *N*=4.

### Metabolites and lipids

In the rhizosphere, the total number of metabolites increased under drought by the end of the 12- week period (moisture*time interaction, p = 0.09; Fig. 5a). The total number of metabolites decreased through time in the detritusphere (p = 0.004), though there was no effect of soil moisture (p = 0.2; Fig. 5a). Similar patterns were observed through direct comparisons of unique metabolites in the rhizosphere versus detritusphere (Fig. 5b). Under normal moisture conditions, the number of unique metabolites increased through time in the rhizosphere, and decreased through time in the detritusphere (Fig. 5b). Under drought conditions, the number of unique metabolites in the rhizosphere increased through time, though stayed relatively consistent through time in the detritusphere.

**Figure 5.**
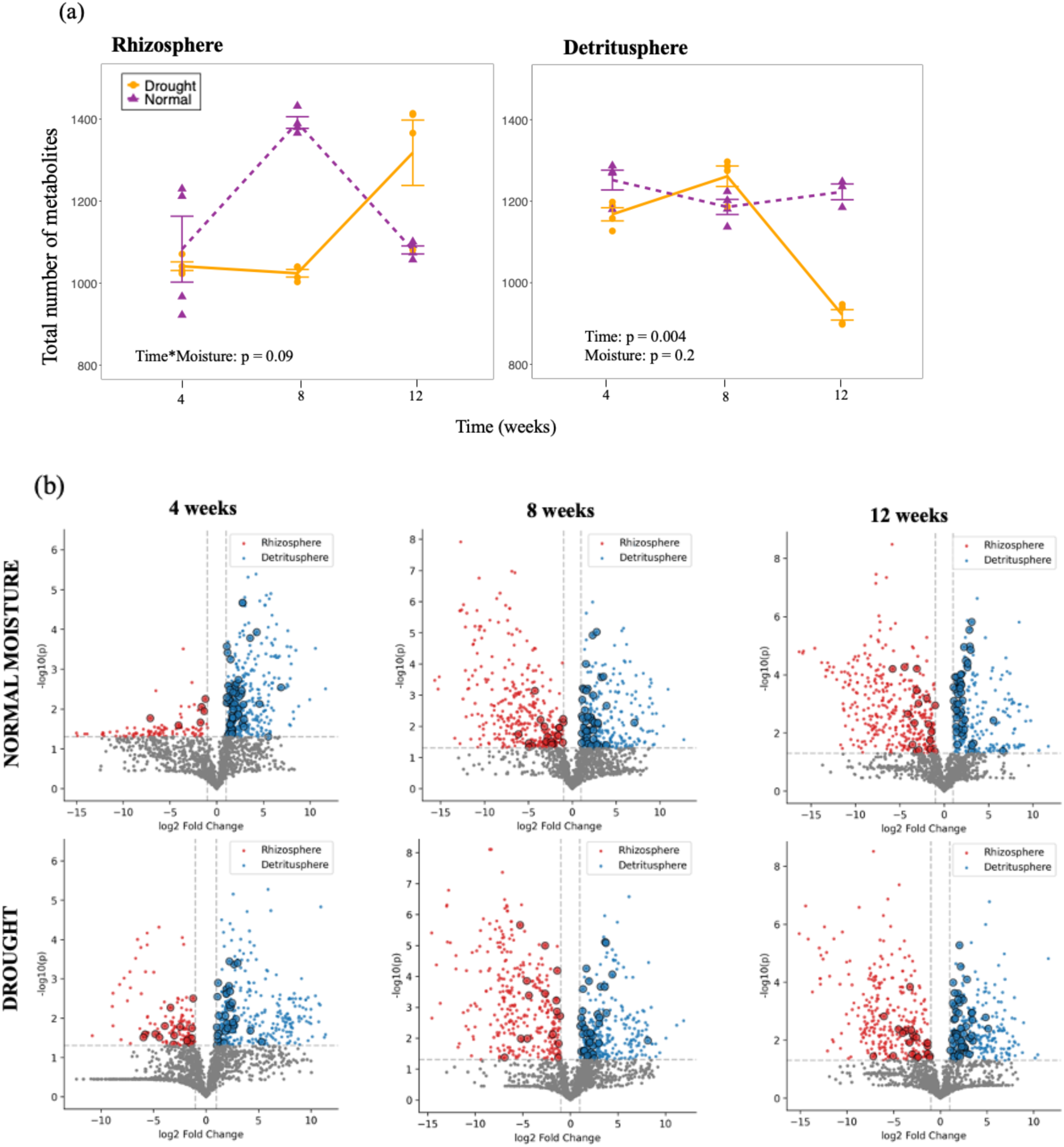
Metabolites in the rhizosphere and detritusphere of a semi-arid grassland soil. (a) Total number of detected metabolites (blank-subtracted) in the rhizosphere and detritusphere. (b) Volcano plots show direct comparisons of metabolites differing significantly in abundance in the rhizosphere (red dots) versus the detritusphere (blue dots) at 4, 8, and 12 weeks (grey dots indicate metabolites found in both habitats; encircled dots indicate an identified metabolite, all others are unidentified metabolites based on current libraries). N = 4.

The total number of unique lipids slightly increased through time in the rhizosphere under normal moisture, though did not change in the detritusphere (Fig. S5). Under drought, the number of unique lipids increased in the detritusphere, but did not change in the rhizosphere (Fig. S5). There were notable differences between the top 25 most abundant lipids in the rhizosphere versus the detritusphere at week 4 and at week 12 (Fig. 6). At week 4, several diacylglycerol lipids and a range of triacylglycerols were more abundant in the detritusphere compared to the rhizosphere (Fig. 6, left panel). At week 12, a range of triacylglycerols and a diacylglycerol were more abundant in the rhizosphere versus the detritusphere, and several glycerolipids, glycerophosphocolines, and glycerophosphoethanolamines were more abundant in the detritusphere (Fig. 6, right panel).

**Figure 6.**
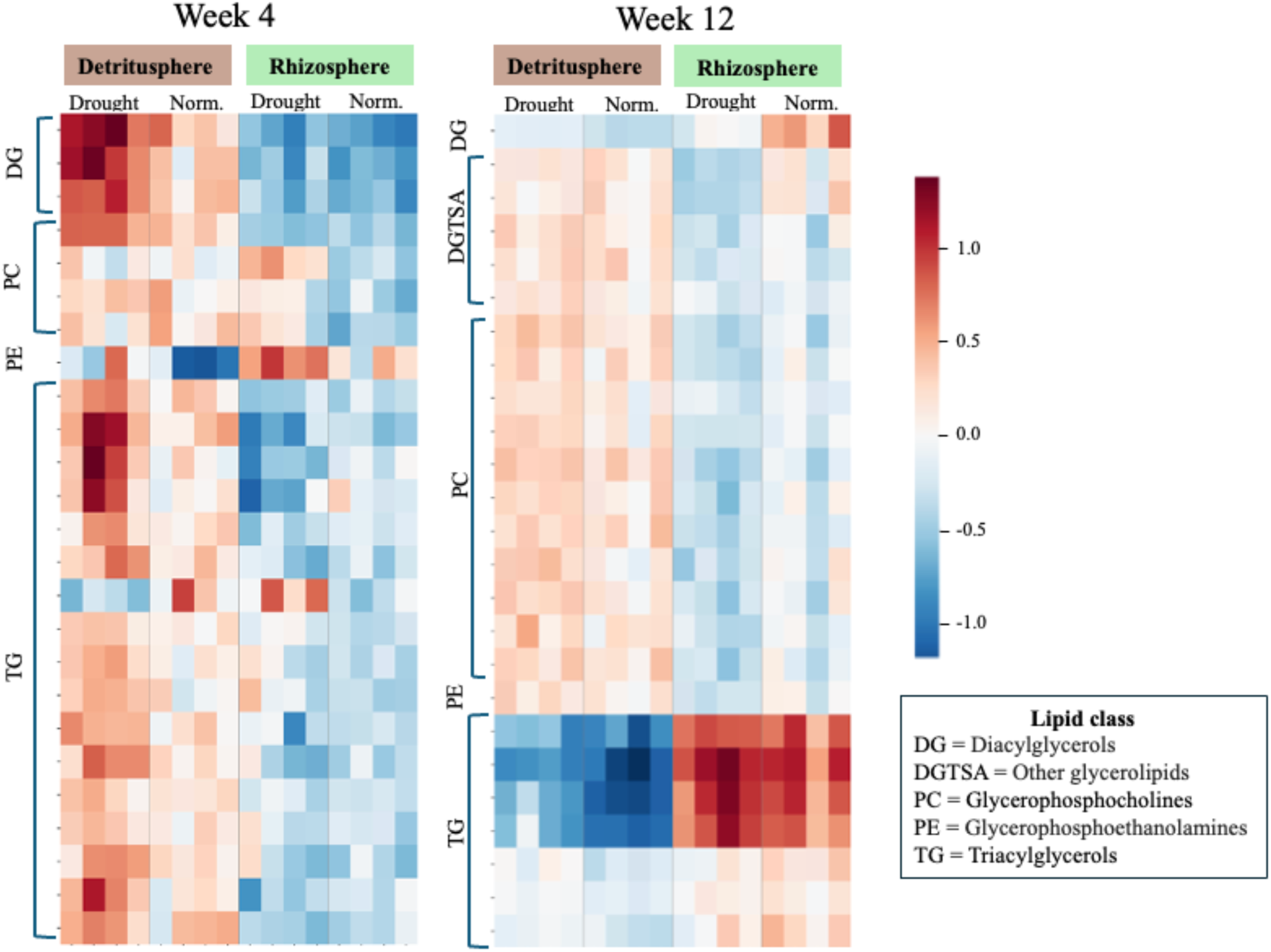
Lipid composition in the detritusphere and rhizosphere during a 12-week ^13^CO_2_- labeling experiment in a semi-arid grassland soil. Heatmaps show the top 25 most significantly different lipids at week 4 (left) and week 12 (right) between the two habitats, with log- normalized relative abundances. Each column represents an individual replicate in each habitat and moisture treatment. *N*=4.

## DISCUSSION

Drought may exert substantial effects on plant growth and microbial activity, but the downstream effects on soil organic carbon have rarely been tested in manipulative studies – especially the formation and persistence of MAOM (Heckman et al. 2023). Here, we evaluated how a spring drought affected MAOM accrual over the 12-week growing period in a semi-arid annual grassland soil, by tracking ^13^C-labeled rhizodeposits of living *Avena barbata* plants into the rhizosphere soil, and ^13^C-labeled *A. barbata* root detritus into detritusphere soil under moisture normal versus droughted conditions. Overall, drought reduced formation of MAOM – both the concentration of new ^13^C-MAOM that formed per gram of rhizosphere or detritusphere soil (µg ^13^C-MAOM g soil^-1^), as well as the total amount of new MAOM formed in the rhizosphere and detritusphere throughout the entire microcosm (µg ^13^C-MAOM microcosm^-1^) (Fig. 1). Drought had the most pronounced effect on rhizosphere ^13^C-MAOM at the end of the growing season (week 12), whereas it had the most pronounced effect on detritusphere ^13^C- MAOM during the early stages of root litter decomposition (i.e., the first 4 weeks). Notably, drought enhanced the persistence of ^13^C-MAOM (i.e., the proportion of ^13^C-MAOM that remained after a 90-day persistence assay) that formed in the detritusphere, though not the rhizosphere (Fig. 2). As we discuss below, the distinct temporal patterns of drought effects in the rhizosphere and detritusphere may be explained by key differences in the quantity and quality of C inputs from living and decaying roots throughout the growing season, and how these C inputs interact with the soil microbial community and mineral matrix to form MAOM.

Drought impacts rhizosphere MAOM formation by affecting the amount and chemical composition of rhizodeposition – with distinct effects throughout the growing season. Plants often release the greatest quantity of rhizodeposits during periods of fast vegetative growth (Eisenhauer et al. 2017; Landl et al. 2021; Gao et al. 2024). Indeed, prior studies with *Avena barbata* have found that its peak growth coincides with the greatest quantity of root exudates released (Zhalnina et al. 2018). In our experiment, peak *A. barbata* aboveground and belowground biomass growth occurred between week 8 and week 12 (Fig. S2). The rhizosphere ^13^C-MAOM pool also showed the greatest increase in size during this same period – both on a per-gram-of soil basis and across the entire microcosm (Fig. 1a-b). It was during this same period of peak plant growth that drought most strongly reduced aboveground biomass (Fig S1a) and belowground root biomass (Fig. S1b) and, in turn, ^13^C-MAOM formation (Fig. 1). Reduced aboveground biomass may have lowered total photosynthate, and thus the rate of rhizodeposition per unit root area (leading to a smaller ^13^C-MAOM pool per g of rhizosphere soil; Fig 1a), while reduced belowground biomass further decreased total C inputs into the soil, diminishing total rhizosphere ^13^C-MAOM throughout the whole microcosm (Fig. 1b). In the field, decreased belowground root biomass should also reduce the total amount of detritusphere C entering the soil profile, but in our experiment we standardized the amount of ^13^C-root detritus entering the detritusphere so we could not detect such an effect. Reduced formation of detritusphere ^13^C- MAOM under drought was thus not the result of reduced initial ^13^C-inputs. Instead, our findings likely reflect the delayed decomposition of ^13^C-root detritus and reduced movement of dissolved OM compounds through the soil matrix under moisture-limited conditions. These factors would affect the timing and transport of the different C compounds that are typically released during the early stages of root litter decay.

Formation of MAOM in both the rhizosphere and detritusphere is affected not only by the total quantity of C inputs, but also by their chemical composition, which influence their direct interactions with the mineral matrix, the rate at which they are decomposed by microbes, and how efficiently they are assimilated into microbial biomass and transformed to MAOM (Oldfield et al. 2018; Chari and Taylor 2022). In the detritusphere, the lowest molecular weight and most soluble root compounds are typically released first during root decomposition (Poll et al. 2008); these compounds are thought to form MAOM most efficiently (Cotrufo et al. 2015; Haddix et al. 2016). Later in the decomposition process, less soluble, more complex polymeric C compounds remain, which are thought to form MAOM less efficiently than simpler compounds (Cotrufo et al. 2015). In support of this pattern, we found that as root litter decayed – observed as a decreasing pool of detritusphere ^13^C-LF over time (Fig. S3) – the average mass of OM compounds increased through time in the detritusphere, and were also higher under drought conditions (Fig. 4). Moreover, protein-like compounds and carbohydrate-like compounds – two compounds classes that are representative of simpler, more soluble compounds – decreased through time in the detritusphere, and also were also more abundant under drought conditions (i.e. due to slower decomposition under drought) (Fig. 4).

The number of metabolites decreased through time in the detritusphere – both the total number of metabolites (Fig. 5a) and the number of metabolites unique to each habitat (Fig. 5b). Such a decrease may suggest that certain metabolites – perhaps smaller, more easily assimilable compounds – preferentially form MAOM early during root decomposition either through the microbial pathway or the direct sorption pathway, and so fewer metabolites remain at the end of the growing season. Moreover, microbial community-level growth rate – which is influenced both by the amount and quality of C input – decreased through time in the detritusphere and was reduced by drought (Fig. 3b). Put together, our results suggest that during the early stages of root litter decomposition, there was a greater proportion of lower molecular weight compounds, a greater total quantity of ^13^C-root detritus (^13^C-LF), a greater number of total metabolites, and the highest microbial community-level growth rate. Drought also had the strongest effect at week 4 on ^13^C-MAOM formation, likely due to delaying the release and transport of lower mass compounds released during the early stages of litter decay that microbes readily consume and that form MAOM most efficiently. By week 12, as less ^13^C-root litter remained, the residual root detritus likely contained larger, more complex compounds (Fig. 4h), and was associated with a lower microbial community-level growth rate (Fig. 3) and fewer total metabolites (Fig. 5a).

We observed opposing temporal patterns in the rhizosphere compared to the detritusphere for the mean mass and composition of OM compounds (Fig. 4), microbial community-level growth rate (Fig. 3) and the number of unique metabolites (Fig 5). Amino sugar-like compounds, protein-like compounds, and carbohydrate-like compounds all increased through time in the rhizosphere – indicating an increase in the amount of low molecular weight compounds (and hence, higher biochemical quality and potentially more microbially assimilable compounds) throughout the growing season (Fig. 4). The average mass of OM compounds in the rhizosphere decreased through time and were lower under drought conditions (Fig. 4d). The total number of unique metabolites in the rhizosphere increased over time (Fig. 5). The ^13^C-LF in the rhizosphere – a combination of fine roots, mycorrhizal inputs, border cells, and other inputs – increased through time (Fig. S3a). Rhizosphere microbial community-level growth rate also increased through time, and was most reduced by drought at week 12 (Fig. 3) – mirroring the pattern observed with the rhizosphere ^13^C-MAOM pool (Fig. 1). Put together, both the quantity and biochemical quality of ^13^C-rhizodeposition increased over the growing season – stimulating greater microbial community-level growth rate and likely greater microbial transformation of rhizodeposition to MAOM (Teixeira et al. 2024; Sokol et al. 2024).

There were notable differences in the lipid profile in the rhizosphere and detritusphere, particularly among triacylglycerols and diacylglycerols, glycerolipids, and glycerophosphocholines (Fig. 6). For instance, at week 4, triacylglycerols were more abundant in the detritusphere compared to the rhizosphere, whereas at week 12, triacylglycerols were more abundant in the rhizosphere. Triacylglycerols are storage lipids that are abundant in fungi and the bacterial genus *Actinomyces* – both of which are known to be resilient to drought conditions in soil (Harwood and Russell 1984; Alvarez and Steinbüchel 2002; Neurath et al. 2021). Their greater abundance in the detritusphere at week 4, especially under drought conditions, may indicate a period of active decomposition by these fungi and filamentous bacteria. Their greater abundance in the rhizosphere at week 12 may indicate greater activity of mycorrhizae and other root-associated microbes, as this corresponded with the period of peak rhizodeposition.

A major outstanding question is how the effects of drought on SOM accrual will persist beyond a single growing season to influence ecosystem C dynamics. In the semi-arid annual grasslands of California, living roots transfer C to the rhizosphere during the spring growing season; those living roots then die during the dry season to form a detritusphere, and that root detritus is largely decomposed in the following growing season (Shi et al. 2018; Fossum et al. 2022; Witzgall et al. 2024). We found that while drought reduced total ^13^C-MAOM in the rhizosphere, it did not affect the persistence of ^13^C-MAOM formed under drought versus normal moisture conditions (Fig. 2a). In contrast, drought did enhance the persistence of ^13^C-MAOM formed in the detritusphere compared to normal moisture conditions (Fig. 2b). Thus, it is possible that while reduced root biomass under drought may decrease the amount of detritusphere MAOM, a greater proportion of that MAOM may persist over time. Prior work on MAOM abundance and persistence in the continental U.S. found that in more arid regions, higher molecular weight and more aromatic compounds were more persistent in soils, perhaps because microbes preferentially used smaller, more soluble compounds (Heckman et al. 2023). Indeed, we found that ^13^C-MAOM derived from more complex root detritus was overall more persistent than ^13^C-MAOM formed from rhizodeposition (Fig. 2). This difference may be explained by the rapid formation and turnover of rhizosphere MAOM, driven by the dynamic interactions of OM with mineral surfaces. These rapid ‘on-off’ dynamics likely result in lower relative persistence of rhizosphere-derived MAOM (Neurath et al. 2021; Heckman et al. 2023). In the detritusphere, drought may have both slowed down microbial activity, selected for certain compounds that form strong associations with mineral surfaces (like greater production of extracellular polymeric substances per g of soil), and/or enhanced interactions between complex OM compounds and mineral surfaces, by drawing together mineral particles and soluble compounds more closely together during soil drying (Kemper et al. 1987; Kaiser et al. 2015; Witzgall et al. 2021).

In conclusion, we provide direct evidence that a spring growing season drought reduces the formation of ^13^C-MAOM relative to normal moisture conditions during the spring growing period in a semi-arid grassland soil. The effects of drought on ^13^C-MAOM formation followed contrasting patterns in the rhizosphere versus the detritusphere of *A. barbata* over the 12-week growing season. These distinct dynamics may be explained by differences in the amount and chemical composition of living root versus decaying root inputs, and how they are processed by the microbial community and transformed to MAOM. Notably, we found that drought enhanced the persistence of MAOM formed in the detritusphere, though not the rhizosphere. Future work will be necessary to determine how these responses affect SOM dynamics over multiple growing seasons in the field, the mechanisms underpinning the effect of drought on MAOM persistence, how the timing and severity of drought events may impact MAOM cycling, as well as how MAOM in other ecosystem types may respond in similar or different ways to drought conditions. Such questions are crucial to understand as increasingly dramatic and unpredictable precipitation patterns pose uncertain consequences for the global carbon cycle.

## ACKNOWLEDGEMENTS

We thank Gianna Marschmann, Ella Sieradzki, Rhona Stuart, Erin Nuccio, Gareth Trubl, Cynthia Jeanette-Mancilla, Melanie Rodriguez-Fuentes, Dinesh Adhikari, Laura Adame, Peter Weber, Rachel Neurath, David Sanchez, Nameer Baker, Sarah Roy, Ilexis Chu-Jacoby, Rachel Hestrin, Emily Kline, Christina Fossum, Mengting Yuan, Alexa Nicolas, Anne Kakouridis, Donald Herman, Craig See, and Aaron Chew for assistance with soil collection, greenhouse harvests and lab analyses; Tina Winstrom at the Oxford Tract Greenhouse at UC Berkeley where the experiment was conducted, and John Bailey and Allison Smith at the Hopland Research and Extension Center where soil was collected. We thank Brad Erkkila at the Yale Analytical and Stable Isotope Laboratory and Jamie Brown at Northern Arizona University for help with stable isotope analyses; Jessica Wollard for assistance with DNA extractions, and Christina Ramon for assistance preparing soil samples for isotope analysis. The LLNL Soil Microbiome SFA team provided valuable feedback on experimental design and data interpretation. This research was supported by the U.S. Department of Energy, Office of Biological and Environmental Research, Genomic Science Program ‘Microbes Persist’ Scientific Focus Area (#SCW1632) at Lawrence Livermore National Laboratory (LLNL) and subcontracts to Northern Arizona University and Pacific Northwest National Laboratory (PNNL). Work conducted at LLNL was conducted under the auspices of the US Department of Energy under Contract DE-AC52-07NA27344.

