## Supplemental Information for "Drought reduces formation, but enhances persistence, of mineral-associated organic matter in a grassland soil"

**Figure S1.** Schematic of experimental design of ^13^CO_2_ greenhouse labeling and incubation experiment, reproduced from Sokol et al., (2024). Shown below is the schematic of a single timepoint, which was replicated for all three timepoints (4, 8, 12 weeks). Root input treatments included (1) rhizodeposits, which were tracked into the rhizosphere in microcosms planted with *Avena barbata*, and (2) root detritus, tracked in the detritusphere – which was defined as the mix of *A. barbata* root detritus + soil contained with a 28-μm mesh bag, buried at the center of an unplanted microcosm. Both types of root input treatment were replicated ten times (*n* = 10) under both normal moisture (~16% soil moisture) and drought (~8% soil moisture conditions. Within each root input × moisture treatment combination, we replicated the ^13^C-labeled treatment 6 times (*n* = 6, shown in red), and a natural abundance unlabeled control 4 times (*n* = 4, shown in blue).

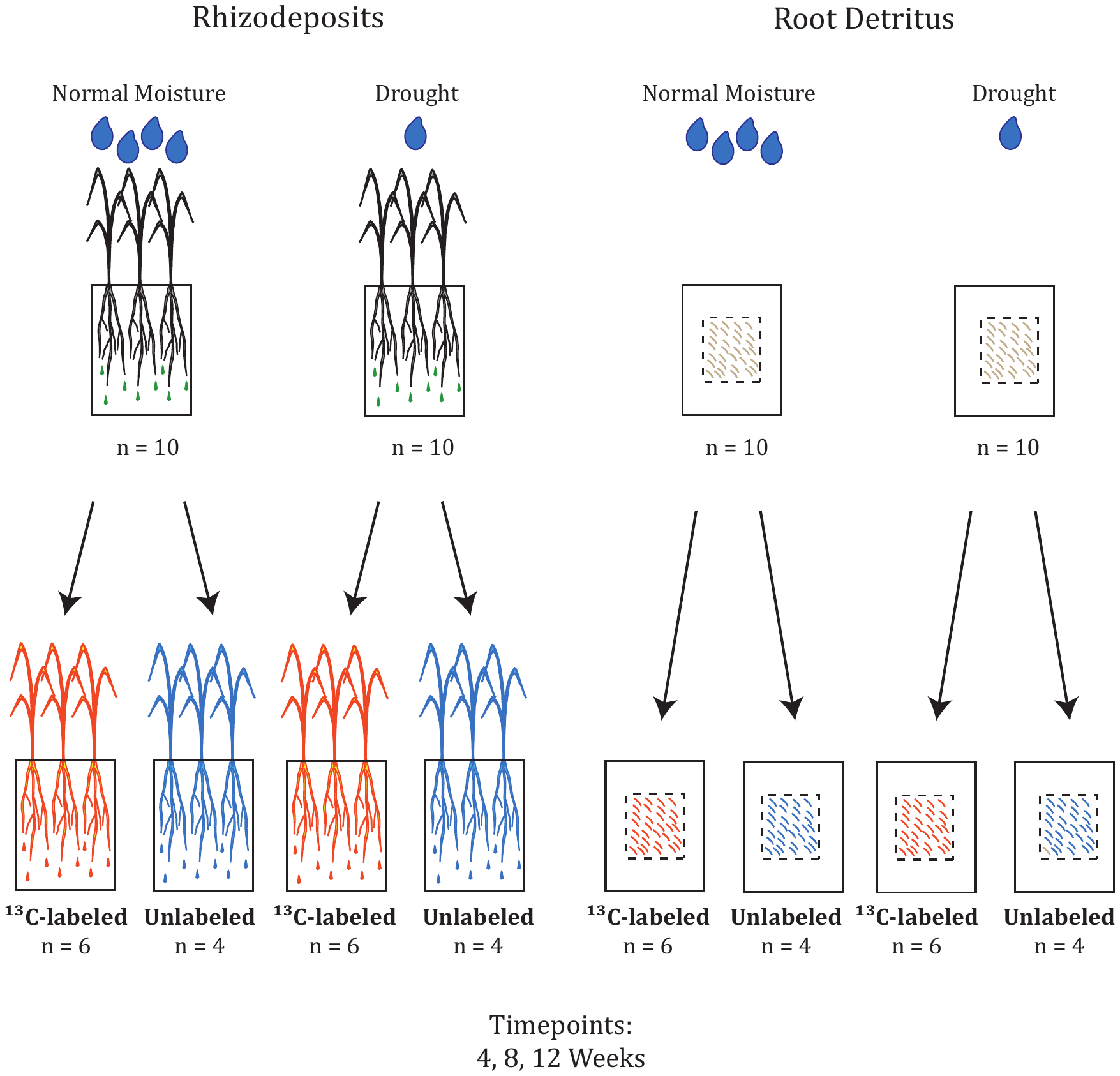

**Fig. S2.** Total aboveground and belowground *A. barbata* biomass formed over a 12-week period in a ^13^CO_2_ greenhouse labeling and moisture manipulation experiment in a semi-arid California annual grassland soil. Shown below is total aboveground biomass (a) and total root biomass (b) in microcosms with living plants (rhizodeposits treatment) under normal moisture (purple) or droughted conditions (orange). Asterisks indicate significant differences between moisture treatments at a given time point (* indicates p<0.05; ** indicates p<0.005). *N* = 6.

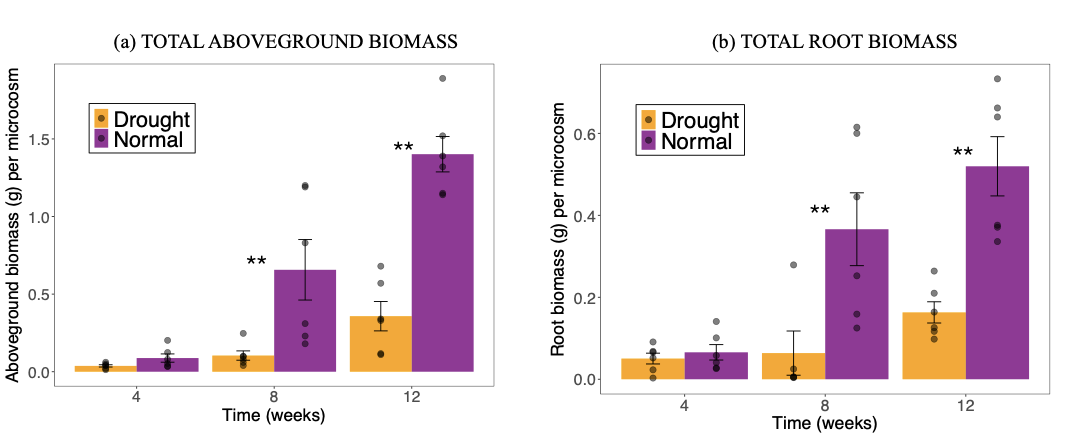

**Fig. S3.** ^13^C-light fraction (^13^C-LF) and ^13^C-coarse heavy associated organic matter (^13^C-CHAOM) formed over a 12-week period in a ^13^CO_2_ greenhouse labeling and moisture manipulation experiment in a semi-arid California annual grassland soil. Shown below is ^13^C-LF and ^13^C-CHAOM formed in the rhizosphere (a,b) and detritusphere (c,d) under normal moisture (purple) or droughted conditions (orange).

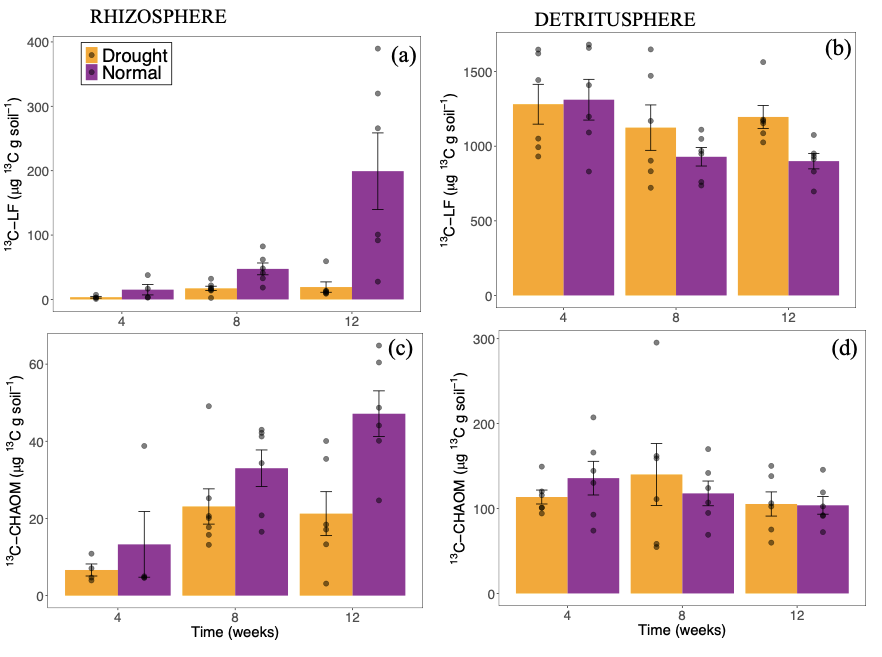

**Fig S5.** Number of unique significant lipids in the rhizosphere versus the detritusphere under normal moisture (left) versus drought conditions (right) during a 12-week growth period/incubation in a semi-arid grassland soil.

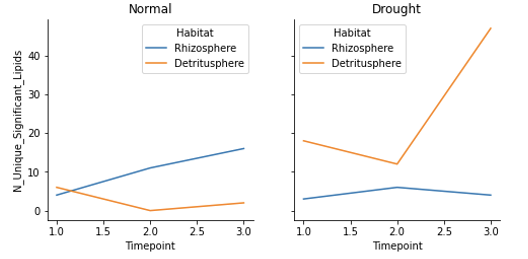

**Supporting Information (Methods):**

*Fourier transform ion cyclotron resonance mass spectrometry (FTICR-MS):*

For each sample, 1g of lyophilized soil was defrosted, added to a clean tube, and extracted with 2 mL milli-Q water (>18.2 MΩ∙cm resistivity). The extraction was performed on 4 replicates (*n* = 4) for each treatment combination at each timepoint. Samples were shaken at 1000 rpm at room temperature for 2 hours using a vortex shaker, centrifuged at 6000 rpm for 5 minutes, and the supernatant was removed. This was repeated with another 2 mL of milli-Q water and supernatants were combined. Solid phase extraction (SPE) was performed on water extracts using Bond Elut PPL cartridges to remove salts and impurities that could interfere with MS analysis (Dittmar et al. 2008). Water extracts were diluted to 5 mL with milli-Q water and adjusted to pH 2 with ~2 µL of 85% H_3_PO_4_ prior to addition to methanol-activated PPL cartridges. Organic matter bound to the cartridges was rinsed with 50 mL of 10 mM HCl, dried with nitrogen and eluted in 1.5 mL of methanol.

Data was acquired using a 7 Tesla Bruker scimaX FTICR-MS operated with quadrupolar (2w) detection and located in the Environmental Molecular Science Laboratory at PNNL (Richland, WA, USA). External calibration was performed using sodium trifluoracetic acid followed by shimming of the analyzer cell to optimize spectra for peak intensity, shape and resolution over the *m/z* range 200-1000. Extracts were direct infused into the electrospray source in negative ion mode at a voltage of +4 kV, dry gas temperature of 200°C and flow of 4 L/min, in randomized order using a custom automated cart. Ion accumulation time was set to 10 ms and three-hundred 8MW time domain transients (2. 1s duration) were coadded, yielding an estimated average mass resolving power of ~680K at *m/z* 400. Internal recalibration of each spectrum was performed with calibration lists of standard OM components. Peak picking was performed in DataAnalysis (v5.0, Bruker Daltonics) using a signal to noise ratio (S/N) threshold of 7, a relative intensity threshold of 0.01 and an absolute intensity threshold of 100,000.

Peak lists were exported and *Formultitude* software (Tolić et al. 2017) was used to align spectra within a 0.5 ppm threshold and assign formulas using N<=2, S=0 and P=0, and <0.5 ppm error for high confidence assignments (mean error < 0.2 ppm). Elemental ratios were calculated from molecular formulas assigned for each peak and averaged for each sample. Molecular class distributions were based on percentage of formulas with elemental ratios within the H/C and O/C ranges for major compound classes (Kim et al. 2003; Tolić et al. 2017) present in each sample. Elemental classifications are putative yet provide a useful indication of the major chemical compound classes and allow for broad comparisons of formula types detected in each sample treatment.

*Metabolomics and Lipidomics:*  Following the water extraction of the soil samples, a modified Folch type extraction was performed to yield polar and non-polar liquid fractions containing ‘metabolite’ and ‘lipid’ components. Ice cold chloroform and methanol (2:1, total 6 mL) were added to the soil, which was vortexed after each addition. Water was added (0.25 mL) and the samples were shaken for 1hr at 1000 RPM. Another 1.25 mL of water was added for a final solvent ratio of 8:4:3 (chloroform:methanol:water), and samples were gently shaken before incubating at 4°C overnight. Samples were then centrifuged at 6000 RPM for 5 minutes, yielding two layers. The layers were separately removed; an aliquot of the bottom (non-polar) fraction was re-added to the top (polar) fraction prior to drying down for subsequent metabolomics analysis, the remaining bottom fraction was dried for lipidomic analysis. Dried samples were stored at –80C prior to analysis.

For metabolomics sample preparation, samples were re-suspended in methanol from the water-extracted SOM SPE eluent at a ratio of 4:1 MeOH:H2O (100 uL) prior to injection (5-10 uL) on the column. LC was performed with a Waters H Class UHPLC. Both a reverse phase and HILIC separation were employed. For reverse phase, a C18 column (Thermo Hypersil Gold, 2.1x150mm, 3um) at 40°C was used with a flow rate of 0.4ml/min and mobile phases of A) 0.1% formic acid in H2O and B) 0.1% formic acid in acetonitrile, with a gradient program of: 0-2 minutes 10% B, 2-11 minutes ramp to 90% B, 11-12 min 90% B, 12-12.5 min increase flow to 0.5ml/min at 90% B, 12.5-13.5 min ramp to 10% B, 13.5-14 min hold 10% B, 14-14.5 min decrease flow to 0.4ml/min, 14.5-16 min hold at 10% B. HILIC separation was performed with a ACQUITY UHPLC BEH HILIC column (2.1x100mm, 1.7um) at 50°C with flow rate of 0.3ml/min and mobile phases of A) 5:95% MeCN:NH4OAc (10mM) in H2O, and B) 100% MeCN with 0.05% NH4OAc. The gradient program was as follows; start 5% A, 0-6 min ramp to 63% A, 6-7 min hold at 63% A, 7-7.1 min ramp to 5% A, 7.1-7.2 min hold 5% A, increase flow to 0.5 ml/min, 7.2-9.5 min hold at 5% A, 9.5-9.7 min hold at 5% A and decrease flow to 0.3 ml/min, 9.7-12 min hold at 5% A.

Metabolomic mass spectrometry measurements were made separately in both positive and negative ion modes with Thermo Q-Exactive Plus (HILIC) or HF-X (reverse phase) Orbitrap mass spectrometers Reverse phase data was acquired with HESI source settings of 3.7/3.0kV high voltage (positive/negative mode), capillary temperature 325°C, sheath gas 35, aux gas 13, and spare gas 3 units, with 30°C probe heating and an S-Lens RF of 50%. HILIC HESI source settings differed with 3.4kV in negative mode, capillary temperature of 320°C and probe heating of 350°C. Reverse phase data dependent experiments were performed with MS1 resolution 240k, AGC target 3e6, max IT 200ms, m/z 80-800, with ddMS2 resolution o 15k performed on top 12 targets with isolation width 0.4 m/z, detection range m/z 200-2000, NCE of 20,30,40 and dynamic exclusion o 30 seconds. HILIC data dependent experiments were performed with MS1 resolution 140k, AGC target 3e6 and max IT 20ms, m/z 80-800, with ddMS2 resolution of 17.5k performed on top 8 targets with isolation with 0.4 m/z, NCE of 20,30,40 and dynamic exclusion of 15 seconds.

*Lipidomics:* Total Lipid Extracts (TLEs) in this study were analyzed by reverse-phase LC-ESI-MS/MS using a Thermo Scientific Vanquish Flex UHPLC system (Thermo Scientific, San Jose, CA) coupled with a Velos Pro Orbitrap mass spectrometer (Thermo Scientific, San Jose, CA). TLEs were stored in 2:1 chloroform:methanol and evaporated, then reconstituted in 5 µl chloroform and 45 µl of methanol prior to injecting 10 µl onto a Waters column (CSH 3.0 mm x 150 mm x 1.7 µm particle size) maintained at 50ºC. Lipid species were separated using a 21 min gradient elution at a 300 µl/min flow rate. Mobile phases A and B consisted of ACN/H2O (40:60) containing 10 mM ammonium acetate and ACN/IPA (10:90) containing 10 mM ammonium acetate, respectively. The full gradient profile was as follows (min, %B): 0,40; 1,62; 4,66; 9,78; 11,87; 15,99; 21,99; 21.1,40. The UPLC system used a Thermo HESI source coupled to the mass spectrometer inlet. The MS inlet and HESI source were maintained at 350ºC with a spray voltage of 3.5kV and sheath, auxiliary, and sweep gas flows of 45, 30, and 2, respectively. Each TLE was analyzed in both positive and negative ion modes in separate runs. Lipids were fragmented by both HCD (higher-energy collision dissociation) and CID (collision-induced dissociation) using a precursor scan of m/z 200-2000 at a mass resolution of 120k, followed by data-dependent MS/MS of the top 4 ions. An isolation width of 2 m/z units and a maximum charge state of 2 were used for CID and HCD scans. Normalized collision energies for CID and HCD were 35 and 30, respectively. CID spectra were acquired in the ion trap using an activation Q value of 0.18, while HCD spectra were acquired in the Orbitrap at a mass resolution of 15k and a first fixed mass of m/z 90.

*Metabolomic data processing:* Confident metabolite identifications were made using Thermo Compound Discoverer 3.3. For RP and HILIC positive and negative mode, spectra were aligned using an adaptive curve with a maximum of 0.3 or 0.6 RT shift respectively and a 3 ppm mass tolerance. Peaks were selected based on a minimum intensity of 1e6 and a chromatographic S/N of 3. Detected features were grouped based on a mass tolerance of 3ppm and a RT tolerance of 0.25. Features were then filtered out depending on the number of samples. Compounds were assigned based on isotopic pattern, RT, MS1, and/or MS2. All identifications and integrated peaks are manually validated and exported for statistical analysis. RT matching is performed on in-house libraries, with MS2 matches based on in-house and external libraries.

Lipidomics data processing: In-house developed software LIQUID (Lipid Informed Quantitation and Identification) (Kyle et al., 2017) was used for all lipid species identifications. This was achieved by importing LC-MS/MS raw data files and examining the diagnostic ion fragments and associated chain fragments in tandem mass spectra. Mass error of measured precursor, isotopic profile, and extracted ion chromatogram (XIC) were also examined for determining lipid identifications. A target database comprised of identified lipid name, retention time, and observed M/Z was used to identify features across all LC-MS/MS runs by aligning and gap-filling the mass spectrometry data of individual sample runs. MZMine2 (Pluskal et al. 2010) was used to align and gap-fill all datasets by matching unidentified features to their corresponding identified features. In addition, peak intensities are exported for statistical analysis after all aligned features are manually verified.
